# Enhancement of Prednisolone efficacy and safety in Duchenne muscular dystrophy via neutrophil elastase inhibition

**DOI:** 10.1101/2025.02.26.640472

**Authors:** K.A. Johnson, F.K. Jones, R.S. Ghadiali, G. Lee, L.A. Torre-Healy, M.D. Sweeney, R. Moffitt, K. Mamchaoui, E. Ricci, I. Young, A Pisconti

## Abstract

Duchenne muscular dystrophy (DMD) is a genetic disorder characterized by muscle wasting, and leading to premature death. The pathogenesis of DMD is complex. It is triggered by loss of the cytoskeletal protein dystrophin, which causes muscle fiber instability and damage, followed by chronic inflammation and fibrotic replacement of damaged muscle tissue. Here we investigated the hypothesis that inhibition of neutrophil elastase, which is increased in dystrophic mice and impairs myogenesis, could provide a therapeutic effect in DMD. While neutrophil elastase inhibition, via the orally bioavailable compound Alvelestat, enhances myogenesis both *in vitro* and *in vivo* and reduces muscle damage and fibrosis, it does not produce a significant beneficial effect on the functional outcomes. However, when co-administered with the standard of care Prednisolone, Alvelestat dramatically enhances Prednisolone efficacy in a mouse model of DMD. In key functional tests, the performance of mice co-dosed with Alvelestat and Prednisolone is almost three times better than that of mice treated with Prednisolone alone. Additionally, co-administration of Alvelestat and Prednisolone reduces Prednisolone’s negative effects on muscle mass and resistance to fatigue. These findings are consistent with the finding that Prednisolone induces neutrophil elastase expression in patients affected by DMD. Our data support a beneficial role for neutrophil elastase inhibition in a mouse model of DMD that is also treated with Prednisolone, suggesting that a combinatorial therapy for DMD including a corticosteroid and a neutrophil elastase inhibitor could be trialed in humans.

**Graphical Abstract:** 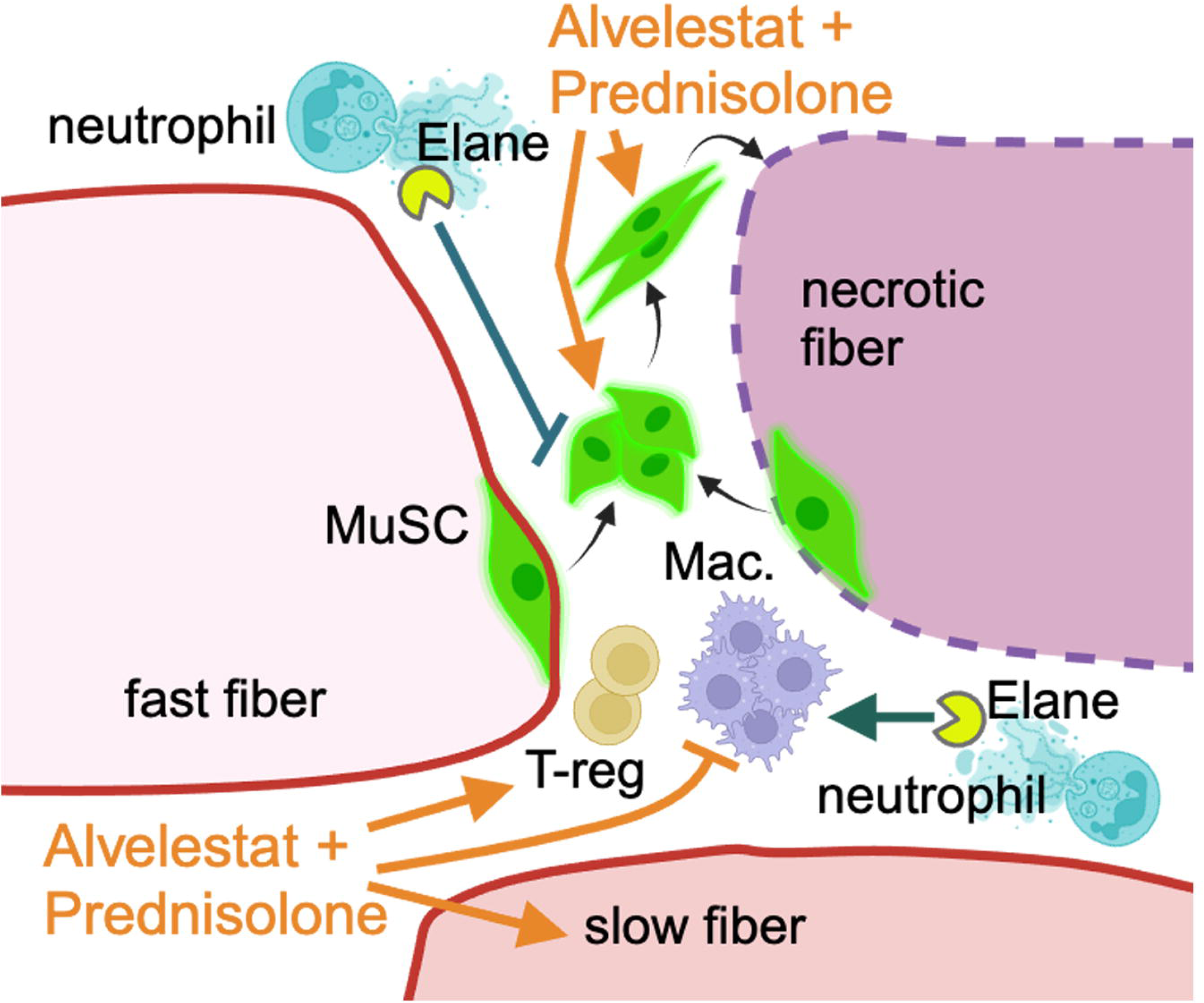

## INTRODUCTION

Duchenne muscular dystrophy (DMD) is a borderline rare genetic disorder and the most common genetic cause of death in boys (1). It is caused by a complete loss of dystrophin, whose full-length isoform is expressed mainly in the muscle tissues, leading to muscle weakness and wasting, and cardiac failure. Corticosteroids, especially prednisone and deflazacort, are still the mainstream standard of care for DMD management, while disease-modifying approaches such as exon skipping, read-through and the recently approved gene therapy, remain only marginally used, both due to their high cost and mutation-dependency (2). Despite their many serious adverse effects, corticosteroids remain the only therapy accessible to all DMD patients, which yield proven benefits, albeit relatively small. The recent approval of the next generation corticosteroid Vamorolone, which yields similar benefits to standard corticosteroids but with reduced side effects (3–5), further encourages the continued use of corticosteroids for the management of DMD and, importantly, their use as benchmark for evaluating the efficacy of new therapeutic approaches.

Neutrophil elastase (ELANE) is one of the three serine proteases released by neutrophils during degranulation and NETosis in response to infection and tissue damage (6). Neutrophils are the first infiltrating immune cells to arrive at the site of injury where they degranulate to release proteases like ELANE that participate in the clearing of the damaged tissue and activation of pro-inflammatory cytokines that attract macrophages (7–9). In response to acute injury, neutrophils are only present at the injury site for approximately two days after which they are cleared via apoptosis and phagocytosis by macrophages. We have recently discovered that the burst of ELANE and other neutrophil serine proteases occurring during the early hours post-injury is essential to promote muscle stem cell activation (Clarke et al., in preparation). However, we have also shown that in the context of chronic injury, such as that associated with DMD, ELANE continues to accumulate over time in a manner that correlates with loss of regeneration capacity (10). *Ex vivo*, chronic exposure to elastase activity causes myoblast death and impairs both proliferation and differentiation.

In this study, we aimed to test the hypothesis that pharmacologic inhibition of ELANE *in vivo* in a mouse model of DMD produces beneficial effects by protecting muscle stem cells and enhancing their regenerative capacity.

## RESULTS

To test our hypothesis that ELANE inhibition is beneficial for DMD, we developed a pre-clinical trial aimed at testing a selective, orally bioavailable ELANE inhibitor in the *mdx^4cv^* mouse model of DMD (Fig. 1). The study was divided in two phases: phase 1 tested three ELANE inhibitors side by side for their potency at inhibiting ELANE *in vitro* and *ex vivo*, and assessed their tissue distribution *in vivo* in *mdx^4cv^* mice (Fig. 1). Upon selecting the lead ELANE inhibitor candidate, phase 2 assessed its efficacy and safety during 12 weeks of treatment of *mdx^4cv^* mice with the lead ELANE inhibitor alongside a standard dose of Prednisolone, which was intended to function as a benchmark. Additionally, we tested the ELANE inhibitor co-administered with Prednisolone (Fig. 1). Control groups were *mdx^4cv^* mice before treatment and at the end of a 12-week-long treatment with placebo (Fig. 1).

**Figure 1.**
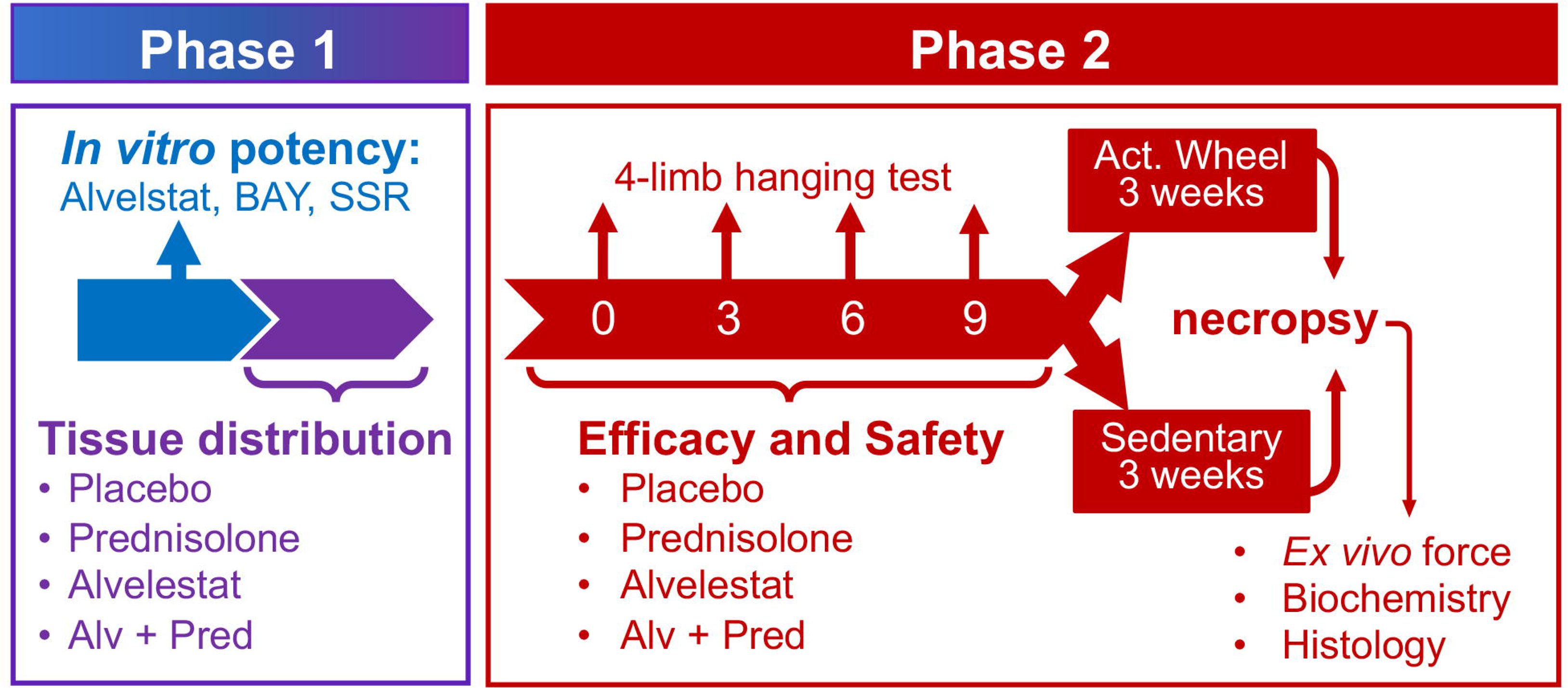
Diagram depicting the experimental design. divided in two phases: phase 1: purple and blue; phase 2: red. The tests carried out during each phase are indicated in the phase-specific color.

### Alvelestat is a potent ELANE inhibitor in vitro and ex vivo, with desirable pharmacokinetics in vivo

Three selective, orally bioavailable ELANE inhibitors were obtained from licensed manufacturers and tested side-by-side for their potency at inhibiting both human and mouse ELANE in an enzymatic activity assay: AZD9668 (a.k.a., and from now on referred to as, Alvelestat), BAY-678 and SSR69071. All three inhibitors showed similar dose-dependent inhibitory activity against human and mouse ELANE, with BAY-678 being slightly less potent and SSR69071 being slightly more potent than the other two inhibitors (Fig. 2A-B). We have previously shown that ELANE induces myoblast death *in vitro*, likely via anoikis(10). To test the potency of the three ELANE inhibitors at preventing myoblast cell death *ex vivo*, we treated human myoblasts derived from a patient affected by DMD(11) with human ELANE (Fig. 2C), and *mdx^4cv^* myoblasts with mouse ELANE (Fig. 2D), in the presence/absence of varying concentrations of the three ELANE inhibitors. The *ex vivo* potency of the three inhibitors against human ELANE, on human DMD myoblasts, reflected the trends already observed *in vitro* when tested against the purified enzyme, with SSR69071 and Alvelestat showing similar potency and BAY-678 showing slightly less potency (Fig. 2C). When tested against mouse ELANE in *mdx^4cv^* myoblast cultures, again BAY-678 showed lower potency at the concentrations tested (Fig. 2D).

**Figure 2.**
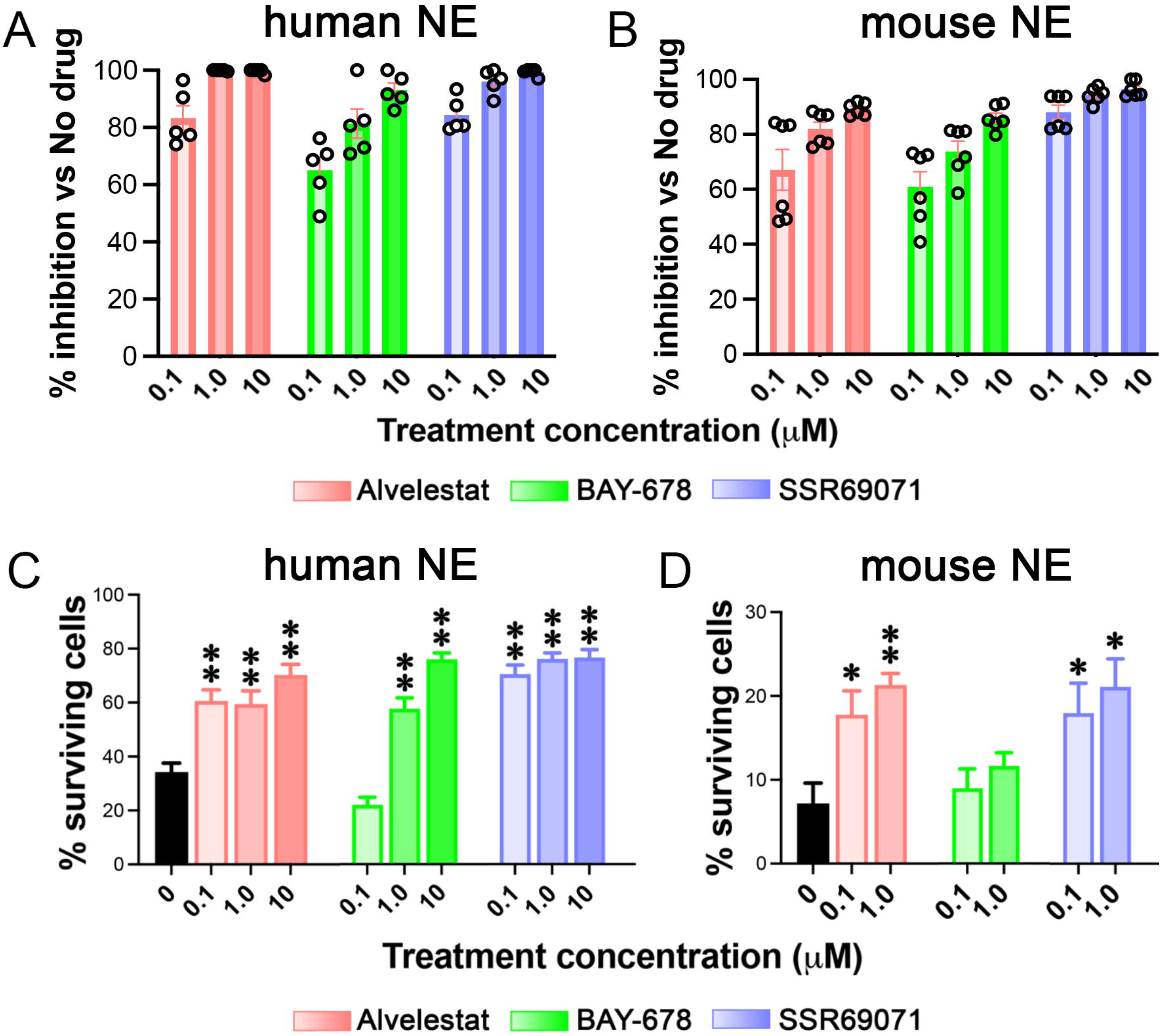
Alvelestat, BAY-678 and SSR69071 have similar inhibitory potency *in vitro* and *ex vivo*. **A-B)** Three increasing concentrations of Alvelestat (pink), BAY-678 (green) and SSR69071 (purple), depicted as increasingly bold colors, were tested side by side for their capacity to inhibit human neutrophil elastase (A) and mouse neutrophil elastase (B) *in vitro* in an enzymatic activity assay. For each condition, three replicates across two independent experiments are plotted as average of the residual activity (ELANE alone without inhibitor was set at 100%) +/- S.E.M. **C-D)** Human myoblasts derived from a patient with DMD (C) and mouse primary myoblasts isolated from *mdx^4cv^* mice (D) were treated with human or mouse, respectively, ELANE in the presence of varying concentrations of Alvelestat (pink), BAY-678 (green) and SSR69071 (purple), depicted as increasingly bold colors. The percentage of survival, compared to control cells that had not received either ELANE or the drugs, is plotted for each condition as the average across three technical replicates and three independent experiments +/- the S.E.M. * = p<0.05 ** = p<0.01, both vs ELANE-treated without drug.

Lastly, we measured the tissue distribution of each ELANE inhibitor *in vivo*, at steady state after a one-week exposure. For each ELANE inhibitor, plasma and muscle (tibialis anterior and diaphragm) levels increased in a dose-dependent manner (Fig. 3). While Alvelestat and BAY-678 were detected at approximately similar levels in plasma and muscle, SSR69071 was detected at much lower levels in all three tissues dosed (Fig. 3).

**Figure 3.**
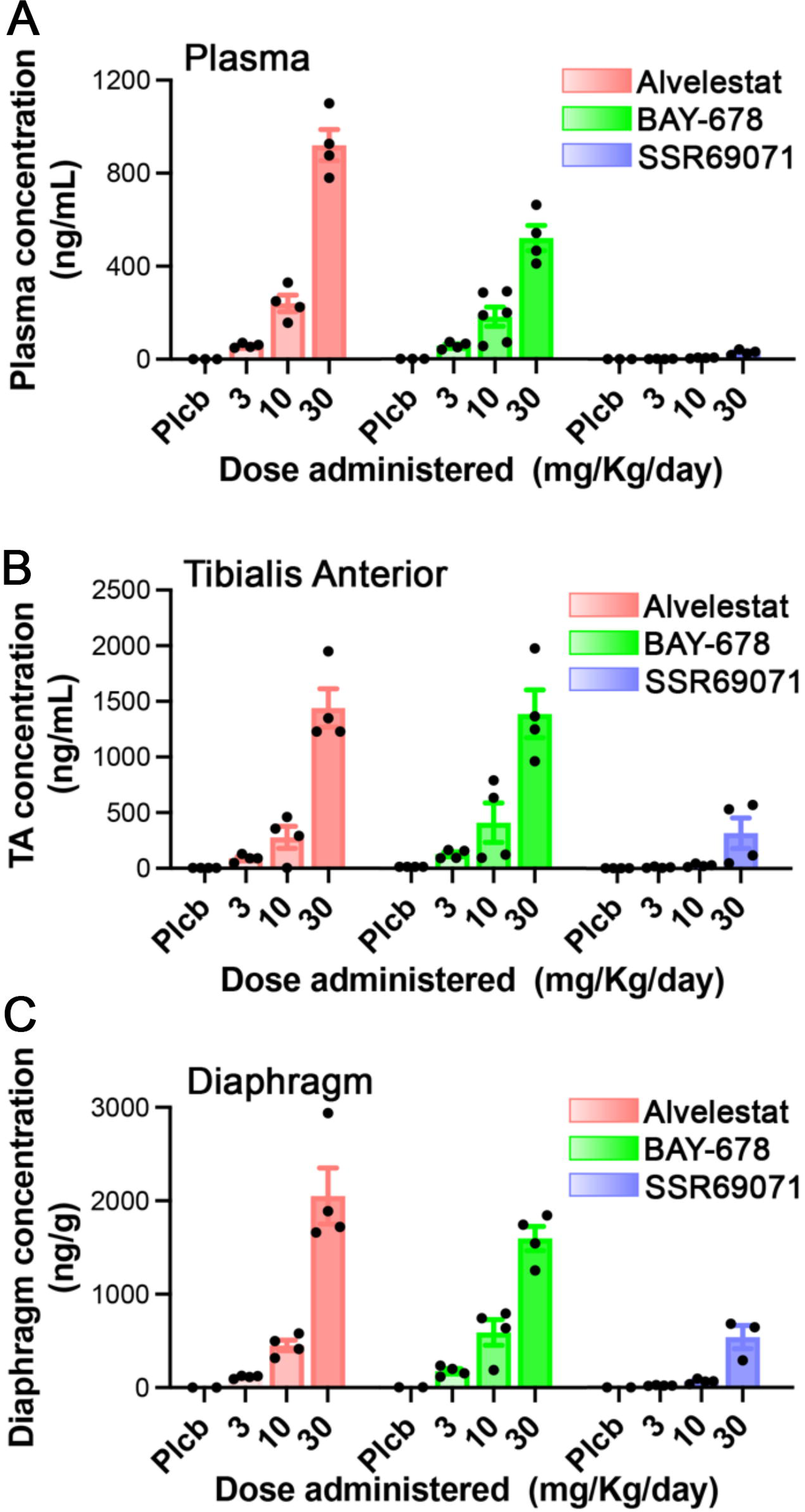
Alvelestat distributes to muscle more effectively than BAY-678 and SSR69071. Three increasing doses (depicted as increasingly bold colors) of Alvelestat (pink), BAY-678 (green) and SSR69071 (purple) were administered to *mdx^4cv^*mice for one week and then the concentration of each drug in (A) plasma, (B) tibialis anterior muscle and, (C) diaphragm muscle measured via mass spectrometry. The average of 4 biological replicates is plotted for each condition +/- the S.E.M.

Although SSR69071 appeared slightly more potent *in vitro* and *ex vivo* compared to the other two compounds, its poor tissue distribution *in vivo* discouraged its further in an efficacy and safety study. Both *in vitro* and *ex vivo*, Alvelestat showed greater potency than BAY-678, whilst their levels in muscle at steady-state exposure were comparable or greater in some cases for Alvelestat. Moreover, Alvelestat has already been in clinical trial and proven highly safe in humans (12–14), while BAY-678 has not. All these data and considerations, taken together, prompted us to abandon both BAY-678 and SSR69071 and continue to study the efficacy of ELANE inhibition in dystrophic mice with Alvelestat.

### Co-administration of Alvelestat and Prednisolone improves Prednisolone’s efficacy in dystrophic mice

Upon identification of Alvelestat as our lead ELANE inhibitor candidate to test *in vivo*, we moved to phase 2 of the study and tested Alvelestat alongside a standard regime of the corticosteroid Prednisolone, and a combinatorial regime including both Prednisolone and Alvelestat (Fig. 1). The primary outcomes of the efficacy study were muscle function and animal fitness – measured as voluntary activity and latency to fall in a four-limb hanging test – while the secondary outcomes were muscle tissue health – assessed via measurement of several biochemical and histological endpoints – and identification of potential molecular mechanisms (Fig. 1).

We used two different functional tests to assess muscle strength and overall fitness: the four-limb latency to fall test, which measures the grip strength of all four limbs, and the voluntary activity wheel, which measures overall mouse fitness (15). The latency to fall test was performed on all mice before the beginning of treatment and every three weeks up to week 9, at which point each treatment cohort was split in two: one half was exposed to an activity wheel and let undertake voluntary physical activity, while the other half was left sedentary to serve as control.

Prednisolone improved limb muscle strength in *mdx^4cv^*mice, as measured by the latency to fall test, during the first weeks of treatment, but this beneficial effect declined later (Fig. 4A and S1A). While Alvelestat treatment of *mdx^4cv^*mice did not lead to a significant improvement of limb muscle strength, surprisingly, the combination of Prednisolone and Alvelestat consistently improved limb muscle strength, with an increase over Prednisolone alone up to approximately 25% at week 9 (Fig. 4A). Importantly, this effect was independent of the body weight (Fig. S1B). The improvement afforded by the addition of Alvelestat to Prednisolone seemed to be mostly due to increased resistance to fatigue, as demonstrated by the fact that the decline in hanging time between the first and third (also last) attempt at hanging within each session, was smaller in mice treated with Alvelestat, either alone or co-administered with Prednisolone, compared with mice treated with either Prednisolone alone or placebo (Fig. 4B).

**Figure 4.**
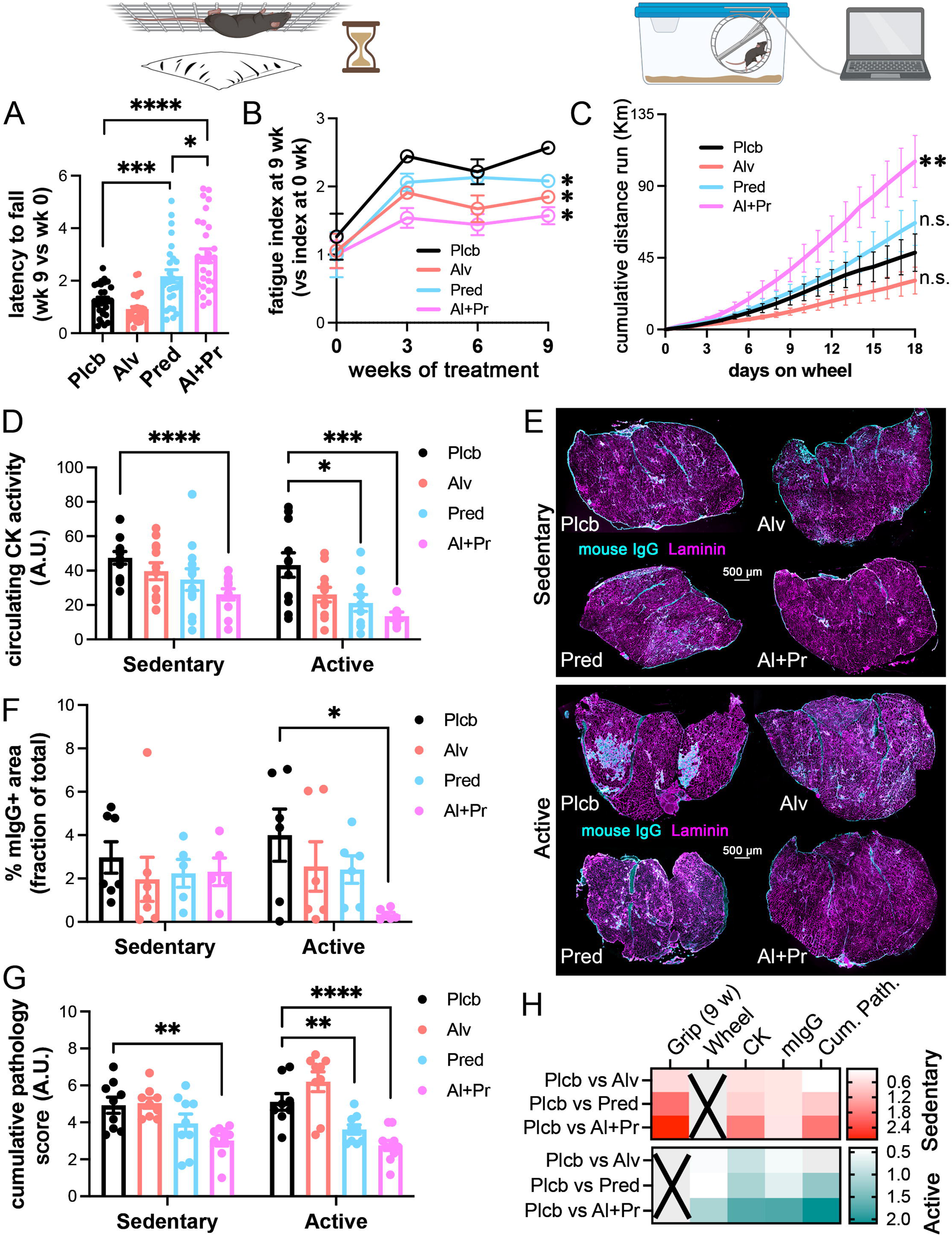
Alvelestat enhances prednisolone efficacy when administered together. **A-B)** *Mdx^4cv^* mice were treated as described in Figure 1 (phase 2) with either placebo (Plcb, black), Alvelestat alone (Alv, pink), Prednisolone alone (Pred, blue) or Alvelestat + Prednisolone (Al+Pr, magenta). The latency to fall in a four-limb hanging test was measured prior to starting treatment (week 0) and then at weeks 3, 6 and 9 into the 12-week program. For each animal the best of six attempts (three per day, 2 days per week) at week 9 was plotted in (A) as the fold change over week 0 (individual points) with the average depicted as bar height and the S.E.M as error bar. In (B) fatigue is plotted at each time point, for each cohort, as the average fatigue index in that week divided by that at week zero. Error bars are S.E.M. N=28 at least, per cohort. **C)** At week 9 each cohort was split in two arms: the animals in the “active” arm were single- house and exposed for three weeks to a voluntary wheel. The cumulative distance run by each animal was calculated and plotted as average per cohort +/- S.E.M. N=12 at least, per cohort. **D)** Levels of creatine kinase (CK) in peripheral blood plotted for each animal in the study (individual points). The averages per cohort are depicted as bar height and the error bars are S.E.M. **E)** Representative images of the gastrocnemius muscle immunostained to detect necrotic fibers (green, mouse IgG), laminin (red) and nuclei (DAPI, blue). **F)** Quantification of (E) across at least 6 animals per cohort. The averages per cohort are depicted as bar height and the error bars are S.E.M. **G)** The triceps muscles were stained with H&E, scored by a veterinarian pathologist and assigned a score 1-4 corresponding to increasing presence of pathological signs. Both triceps were scored for several parameters and for each animal the total of scores of all parameters, averaged across the two triceps, are plotted as individual points. The averages per cohort are depicted as bar height and the error bars are S.E.M. **H)** For several of the outcomes shown in this figure we were able to calculate the effect size (Cohen’s *d*) of Prednisolone and Alvelestat, both alone and in combination.

The positive effect of Alvelestat addition to Prednisolone was even more obvious when overall mouse fitness was assessed as the cumulative physical activity voluntarily undertaken over a period of 18 days (Fig. 4C). The improvement in overall fitness produced by Prednisolone alone was not statistically significant (Fig. 4C) and the effect size was small (Fig. 4H), while the beneficial effect of the Prednisolone and Alvelestat combination was highly statistically significant and nearly three times greater than Prednisolone alone (Fig. 4C and 4H).

We next assessed the extent of myofiber damage by measuring both the circulating levels of creatine kinase activity (Fig. 4D) and the percentage of necrotic myofibers, which uptake mouse immunoglobulins (Fig. 4E-F). In both cases, we observed an improvement in mice treated with Prednisolone, which was further enhanced by the co-administration of Alvelestat, especially in active mice (Fig. 4D-F and H), as previously observed with the functional outcomes (Fig. 4A-C and H).

In addition to mIgG uptake, we performed a comprehensive assessment of muscle pathology on triceps sections stained with hematoxylin and eosin, that included assessment of: (i) inflammation (presence of immune infiltrates), (ii) fiber mineralization, (iii) degeneration (presence of hypercontracted fibers), and (iv) regeneration (presence of myotubes, centrally-nucleated basophilic fibers). We found once again that Prednisolone decreases the cumulative pathology score of these muscles and co-administration of Alvelestat and Prednisolone further decreases the score, especially in animals that undertook voluntary physical activity (Fig. 4G).

In summary, coadministration of Alvelestat and Prednisolone dramatically enhanced the effect size of the beneficial effect of Prednisolone on all the outcomes measured, even though Alvelestat alone was mostly ineffective (Fig. 4H).

### ELANE inhibition rescues Prednisolone-induced susceptibility to myofiber fatigue

To assess whether the increased fitness and resistance to fatigue (Fig. 4A-C) observed in mice treated with the combination of Prednisolone and Alvelestat was due to an intrinsic increase in resistance to myofiber fatigability, we measured the force of diaphragm strips *ex vivo* in response to a tetanic stimulus administered for 30 seconds (Fig. 5A). We chose the diaphragm muscle because is one of the most severely affected in *mdx^4cv^* mice and because it is amenable to *ex vivo* force measurements due to its high surface-to-volume ratio. Muscles of mice treated with Prednisolone rapidly lost force during the tetanic contraction compared with placebo-treated mice (Fig. 5B-C). This negative effect is fully reverted in active mice and partly in sedentary mice by the co-administration of Alvelestat to Prednisolone, with Alvelestat alone improving also force retention, especially in sedentary mice (Fig. 5B and D). The beneficial effect on fatigue conferred by co-administration of Alvelestat and Prednisolone was consistent with the increased resistance measured through the four-limb latency to fall test. It was not mediated by an increase in the force elicited by individual contractile units since the peak specific force of diaphragm strips from mice treated with both Prednisolone and Alvelestat together was, in fact, slightly smaller than that of muscle strips from mice treated with Prednisolone alone (Fig. S2A). Thus, our data suggest that (i) Prednisolone significantly increases muscle-specific force but decreases resistance to fatigue in dystrophic mice; (ii) co-administration of Alvelestat and Prednisolone rescues Prednisolone-induced loss of resistance to fatigue, while maintaining specific force essentially unaltered. This could be explained if the addition of Alvelestat to Prednisolone induced a myofiber type switch towards slower types, and indeed we observed a significant increase in type I and type IIa fibers in mice co-dosed with Prednisolone and Alvelestat at the expense of type IIx and type IIb fibers (Fig. S2B).

**Figure 5.**
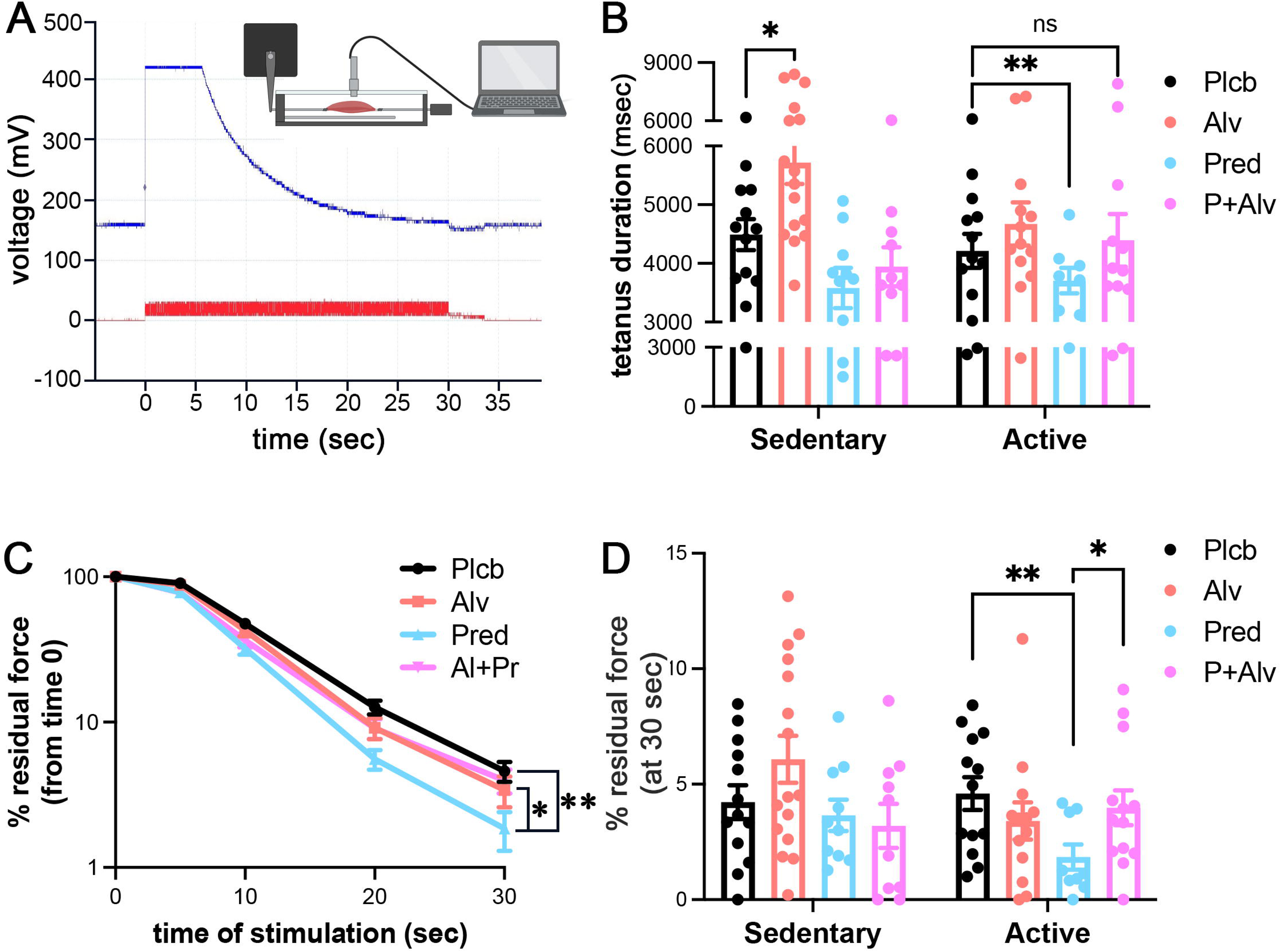
Alvelestat addition rescues Prednisolone-induced myofiber fatigability. **A)** Representative force trace (blue) in response to a tetanic stimulus (red). **B)** Quantification of the duration of the tetanic phase [top flat part of the blue curve in (A)] of the force elicited by a tetanic stimulus across all animals in the study (individual points) sorted based on whether belonging to the sedentary or the active arm in the last 3 weeks of the study. The averages per cohort are depicted as bar height, and the error bars are S.E.M. N=12 at least per cohort. **C-D)** Muscle fatigue is measured as the percentage of residual force, relative to the peak force, during (C) and at the end (D) of the tetanic stimulus. In (C), the averages per cohort are plotted +/- S.E.M.; in (D), the averages per cohort are depicted as bar height, and the error bars are S.E.M. N= 12 at least per cohort.

#### Co-administration of Alvelestat and Prednisolone rescues Prednisolone-induced muscle atrophy and liver toxicity

Long-term treatment with corticosteroids, such as Prednisolone, induces significant muscle atrophy and loss of body weight in dystrophic mice(16, 17). Indeed, we found that after 12 weeks of daily Prednisolone treatment, *mdx^4cv^* mice lose on average over 12% of their body weight while the body weight of mice treated with Alvelestat does not significantly change (Fig. 6A). Moreover, co-administration of Alvelestat to Prednisolone slightly decreases Prednisolone-induced body weight loss, especially in the mouse cohort that remained sedentary (Fig. 6A). It is possible that the effect size of Alvelestat on Prednisolone-induced body weight loss is small due to a general beneficial effect of exercise on body fat. Indeed, mice co-dosed with both Prednisolone and Alvelestat exercise more (Fig. 4C) and therefore are more likely to lose weight simply as an effect of increased physical activity, as observed in age- and sex-matched placebo-treated wild-type mice (Fig. S3). Importantly, when we compare muscle weight across all cohorts, we observe that gentle exercise does indeed promote muscle growth in mice treated with Alvelestat, to the point that Prednisolone-induced muscle loss is almost completely rescued by the addition of Alvelestat in the active cohort (Fig. 6B). To assess whether this is a direct effect of Alvelestat on Prednisolone-induced muscle atrophy, we cultured a myoblast cell line derived from a patient affected by DMD (11) and after inducing differentiation, we treated the myotubes with either Prednisolone alone, Alvelestat alone, or in combination (Fig. 6C). As expected, Prednisolone induced myotube atrophy in a cell-autonomous manner, which was fully rescued by addition of Alvelestat (Fig. 6C).

**Figure 6.**
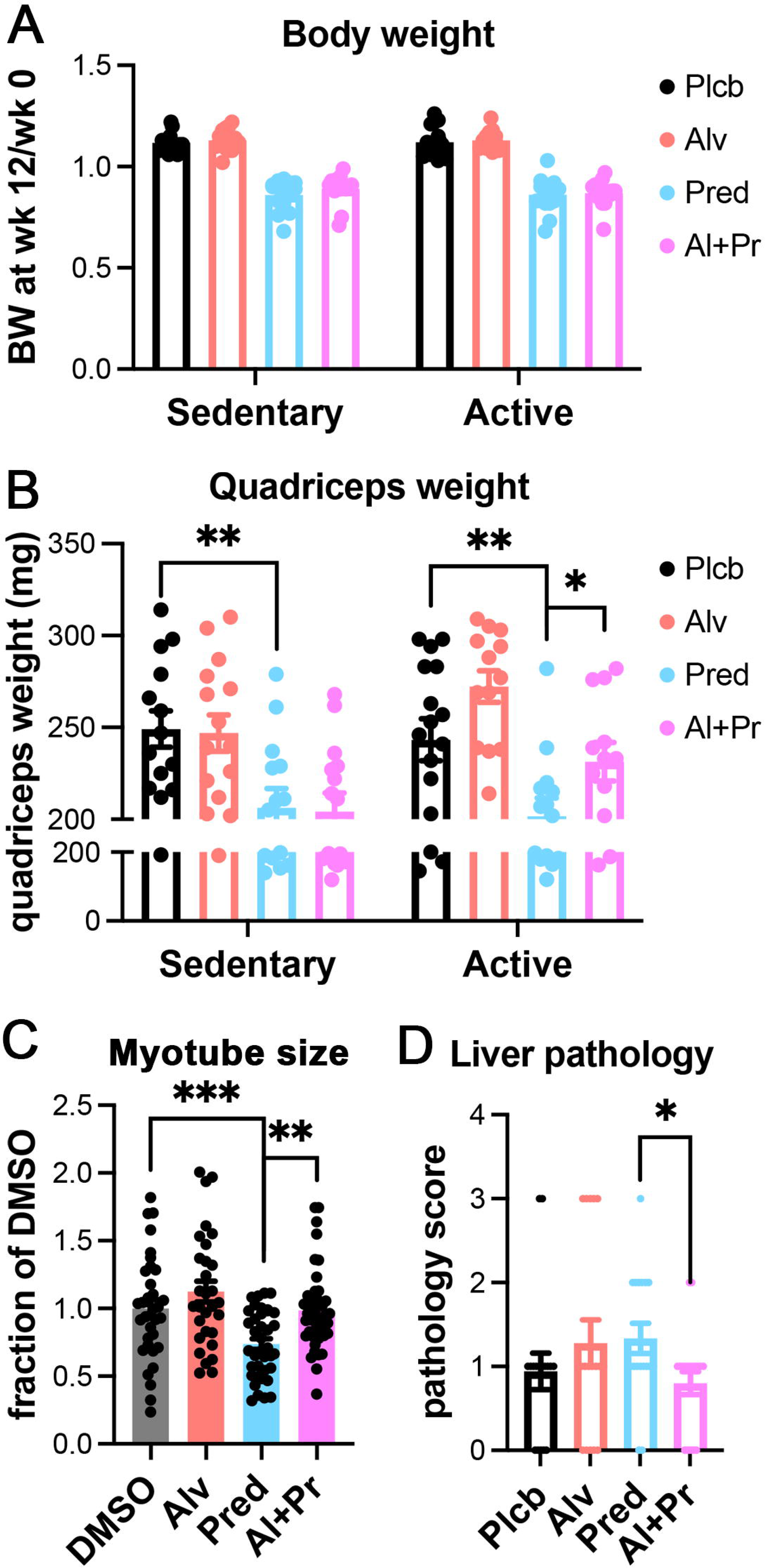
Alvelestat co-administration rescues Prednisolone-induced muscle atrophy in active animals and liver injury. **A)** The body weight of each animal was measured and plotted as fold change at the end of treatment over week 0. Alvelestat did not significantly improve Prednisolone-induced weight loss, but **B)** muscle weight was increased by Alvelestat, both alone and in mice treated with Prednisolone, in the active group. **C)** Myoblasts derived from a patient affected by DMD were differentiated into myotubes then exposed for 2 days to Alvelestat or Prednisolone, alone or in combination, or vehicle (DMSO) as indicated, then fixed, immunostained to detect myosin heavy chain (MF20 clone) and processed to calculate the total myotube area. **D)** Livers were sectioned, stained with H&E, scored by a veterinarian pathologist and assigned a score 1-4 corresponding to increasing presence of pathological signs. Scores for each animal are plotted (individual points). Note: no differences were noted within the same drug cohort between the active and sedentary mice. The averages per cohort are depicted as bar height and the error bars are S.E.M.

Another important side effect of long-term Prednisolone treatment of *mdx^4cv^* mice that emerged from our study is mild liver toxicity, as evidenced by histopathological examination (Fig. 6D). While Alvelestat alone also induced the same extent of mild liver toxicity, the co-administration of Alvelestat and Prednisolone appeared to have surprisingly less liver toxicity than either drug alone (Fig. 6D).

#### Inflammation and fibrosis are differentially regulated by Alvelestat and Prednisolone

Treatment with Prednisolone and Alvelestat should decrease inflammation since: (i) Prednisolone is a potent anti-inflammatory drug (18, 19); (ii) ELANE, which is inhibited by Alvelestat, is a pro-inflammatory protease (7, 8); (iii) mice co-dosed with Prednisolone and Alvelestat show reduced myofiber damage. To test this prediction, we quantified specific and unique markers of immune cells in whole muscle to assess muscle-wide inflammation. This approach is preferred to immunohistochemistry due to the focal nature of muscular dystrophy lesions and is more practical than flow cytometry when analyzing hundreds of animals. As expected, we observed reduced total immune cell burden (CD45, Fig. 7A) in animals treated with Prednisolone. Interestingly, while Alvelestat alone had the opposite effect on CD45, especially in physically active animals (Fig. 7A), co-administration of Alvelestat and Prednisolone further enhanced the anti-inflammatory effect of the latter, similar to the trend observed in the muscle function outcomes as well as in muscle damage and muscle fatigue outcomes (Fig. 4A-C). To gain further insight into this paradoxical effect of Alvelestat on immune cell infiltration into dystrophic muscle, we looked at markers of specific effector cell subpopulations. The vast majority of immune cells infiltrating dystrophic muscles are macrophages (20, 21). When we quantified the general macrophage marker F4/80, we observed a similar trend to that of CD45 (Fig. 7B). Intriguingly, when we quantified the burden of regulatory T lymphocytes (Tregs, FoxP3, Fig. 7C), which have been proposed as key anti-inflammatory cells in DMD (22), we observed an increase in Tregs in mice treated with both Prednisolone and Alvelestat, which was especially noticeable, and statistically significant, in physically active mice (Fig. 7C).

**Figure 7.**
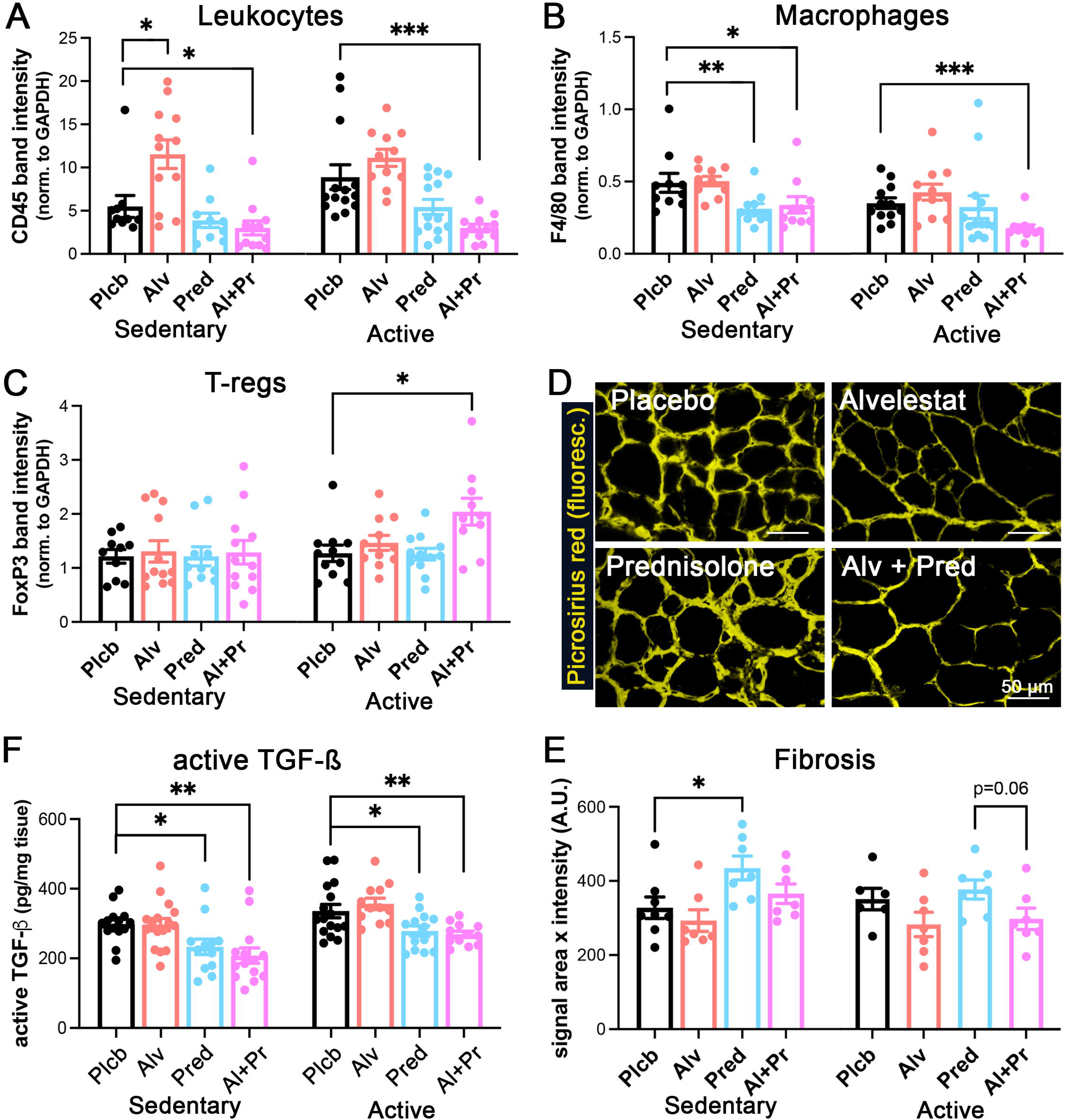
Inflammation and fibrosis are differentially affected by Alvelestat and Prednisolone. **A-C)** Unique markers of immune cell populations – A) CD45 for all leukocytes; B) F4/80 for all macrophages; C) FoxP3 for all T-regs – were quantified in all samples via western blotting. Population- specific band intensity was normalized to GAPDH band intensity for each biological replicate and then all biological replicates were normalized to an internal standard loaded across all gels. These normalized values for each animal are plotted in (A-C) as individual points, sorted based on whether belonging to the sedentary or the active arm in the last 3 weeks of the study. The averages per cohort are depicted as bar height and the error bars are S.E.M. N= 9 at least per cohort. **D)** Representative images of Picrosirius fluorescence displayed in pseudocolor yellow. **E)** Quantification of D) across at least 7 biological replicates per cohort, where the value for each animal is plotted (individual points), and animals are sorted based on whether belonging to the sedentary or the active arm. The averages per cohort are depicted as bar height and the error bars are S.E.M. **F)** ELISA quantification of active TGF-β in muscle extracts. The value for each biological replicate (animal), across all animals in the study is plotted (individual points) sorted based on whether belonging to the sedentary or the active arm. The averages per cohort are depicted as bar height and the error bars are S.E.M. * = p<0.05, ** = p<0.01, *** = p<0.001, in the indicated comparisons.

In muscular dystrophy, chronic inflammation leads to impaired extracellular matrix (ECM) remodeling and chronic accumulation of ECM (fibrosis). We used picrosirius red staining, which detects fibrillar collagen, to assess the extent of fibrosis in the severely affected gastrocnemius muscle and found that, despite the observed decrease in the overall burden of immune cells, mice treated with Prednisolone showed increased fibrosis compared to placebo-treated and Alvelestat-treated mice (Fig. 7D-E). This negative effect of Prednisolone was rescued by the co-administration of Alvelestat (Fig. 7E) and was independent of the effect of either drug on the expression level of collagen genes that make up fibrillar collagen (Fig. S4), nor did it correlate with the effect of Prednisolone and/or Alvelestat on the levels of active TGF-β in muscle (Fig. 7F), which instead correlated with the macrophage burden.

#### Alvelestat does not autonomously affect the transcriptional activity of Prednisolone

Corticosteroids are potent transcriptional regulators and thus, we hypothesized that the beneficial effects observed when Alvelestat is co-administered with Prednisolone are mediated by changes in the transcriptional activity of Prednisolone. However, RNA-sequencing of quadriceps muscles revealed that Alvelestat had no direct effect on the transcriptional activity of cells present in the muscle tissue, as the transcriptome of muscles from mice treated with Alvelestat alone had no differentially expressed genes when compared with placebo (Fig. S5A). More importantly, Alvelestat does not directly alter the effects of Prednisolone on transcription, since the transcriptome of mice co-dosed with Alvelestat and Prednisolone was not different from that of mice that had received Prednisolone alone (Fig. S5B). However, when either Prednisolone alone or the Prednisolone + Alvelestat combination were each one compared to Placebo, a significant set of genes were found differentially expressed uniquely in mice treated with either regimen; namely: 1,486 genes were found differentially expressed only in mice co-dosed with Alvelestat and Prednisolone compared to placebo and 608 genes were found differentially expressed only in mice treated with Prednisolone alone compared to placebo (Fig. 8A). This observation suggests that the co-administration of Alvelestat and Prednisolone somewhat indirectly alters the transcriptional outcome of Prednisolone treatment (Fig. 8A). Functional analysis of the genes that were uniquely altered by Prednisolone alone included *Protein transport* and *Transcriptional regulation*, as to be expected (Fig. 8B). In contrast, the genes whose expression was uniquely altered by co-administration of Prednisolone and Alvelestat affected also *Inflammation* (Fig. 8C, consistent with data presented in figure 7) and metabolic pathways such as *Lipid metabolism* and *Transition between fast and slow fiber* (Fig. 8C-D) consistent with fast-to slow fiber switch presented above in Figure S2B.

**Figure 8.**
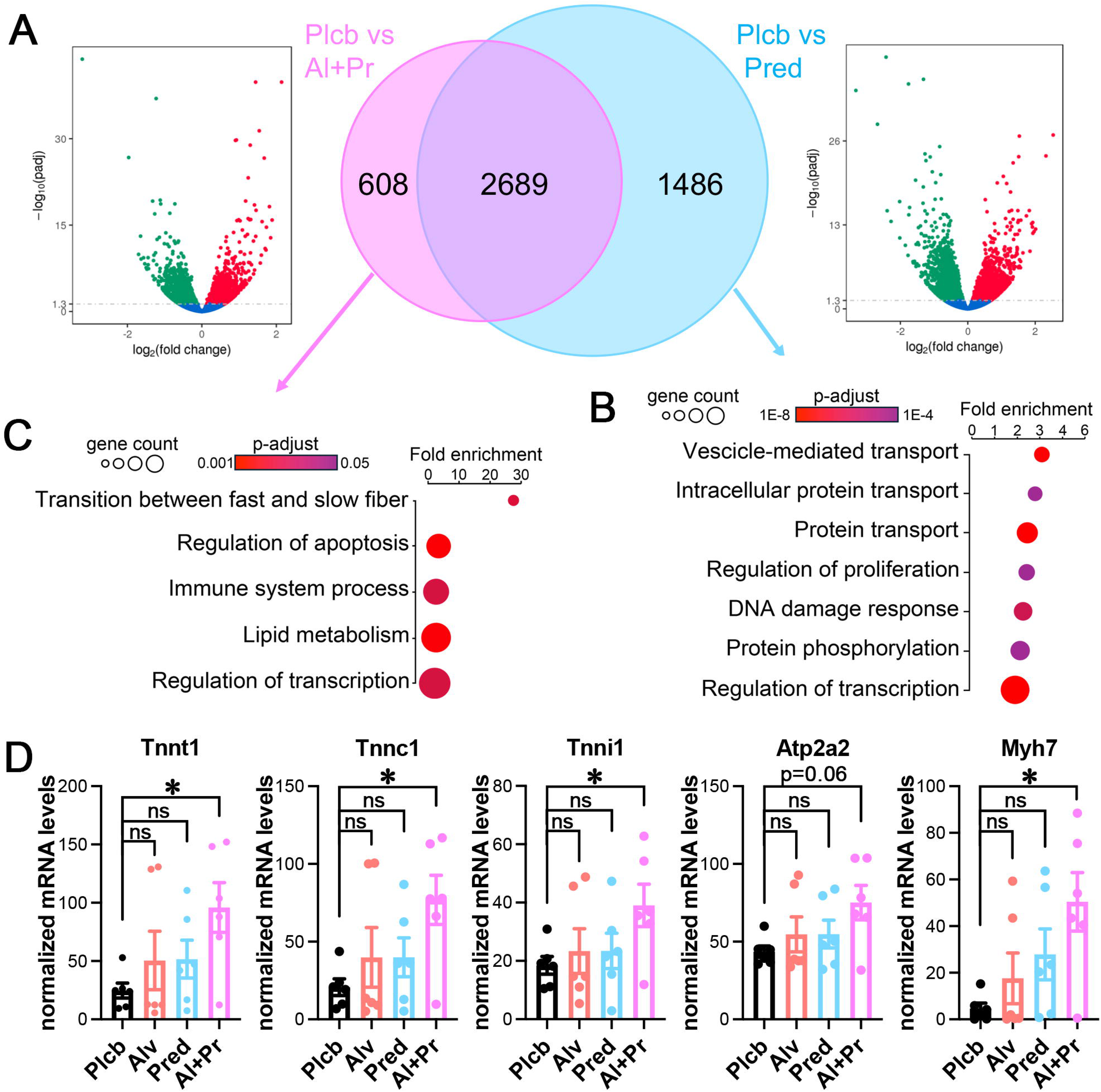
Alvelestat indirectly affects the outcome of the transcriptional activity of Prednisolone. **A)** Volcano plots of the genes differentially expressed in mice treated with Alvelestat + Prednisolone (left) or with Prednisolone alone (right), both compared with mice treated with placebo. Center: Venn diagram depicting the total number of genes differentially expressed in mice treated with Alvelestat + Prednisolone (magenta, 608 unique) or with Prednisolone alone (blue, 1486 unique), both compared with mice treated with Placebo. Gene expression was measured by RNA-sequencing of quadriceps muscles. N=6 per cohort. **B)** GO Biological Processes to which genes differentially expressed in mice treated with Prednisolone alone mapped. **C)** GO Biological Processes to which genes differentially expressed in mice treated with Prednisolone + Alvelestat mapped. **D)** Expression levels of differentially expressed genes within the “*Transition between fast and slow fiber*” GO-BP category, showing a switch from fast to slow type in mice co-dosed with both Alvelestat and Prednisolone.

#### Alvelestat enhances myogenesis *in vivo* and *in vitro*

An important aspect of DMD pathology is the impairment in muscle regeneration. We were prompted to study the effect of ELANE inhibition on muscular dystrophy *in vivo* by our previous finding that chronic exposure to elastase activity impairs myoblast proliferation, survival and differentiation *in vitro* (23). Thus, we assessed the extent of regeneration in the muscles of mice treated with placebo, Prednisolone, Alvelestat or their combination, using a range of biochemical assays and parameters. Firstly, we determined the levels of MyoD1 and embryonic myosin heavy chain (eMyHC) by western blotting, as a measure of muscle stem cell commitment to the myogenic lineage (MyoD1) and of terminal myogenic differentiation (eMyHC). We developed a *Normalized Regeneration Score* by summing the placebo-normalized MyoD1 levels in each treatment to the placebo-normalized eMyHC levels in each treatment, and then dividing the sum by the average CK levels for each cohort. Dividing by the CK levels is a way to normalize the extent of regeneration to the extent of degeneration and, thus, quantify the effect of Alvelestat, Prednisolone and their combination on promoting regeneration, decoupled from their effect on preventing degeneration. An increase in regeneration would be favorable only in the absence of an increase in degeneration, hence the importance of decoupling the two effects. In mice kept sedentary, the Normalized Regeneration Score was not altered by either Alvelestat or Prednisolone, but was dramatically increased by co-administration of Alvelestat and Prednisolone (Fig. 9A). When regeneration was stimulated via physical activity, we observed an increase in Normalized Regeneration Score in mice that received Alvelestat or Prednisolone alone and again a dramatic increase in Normalized Regeneration Score when Alvelestat and Prednisolone were co-administered (Fig. 9A). These trends were largely confirmed when regeneration was assessed as the fraction of centrally nucleated myofibers, again normalized by the levels of CK to account for the confounding factor generated by changes in the extent of muscle damage (Fig. 9B). Crucially, the abundance of muscle stem cells in animals treated with Alvelestat, both alone and especially in combination with Prednisolone, was likely increased compared to both placebo and Prednisolone alone, since the mRNA levels of the muscle stem cell unique marker Pax7 were increased (Fig. (9C).

**Figure 9.**
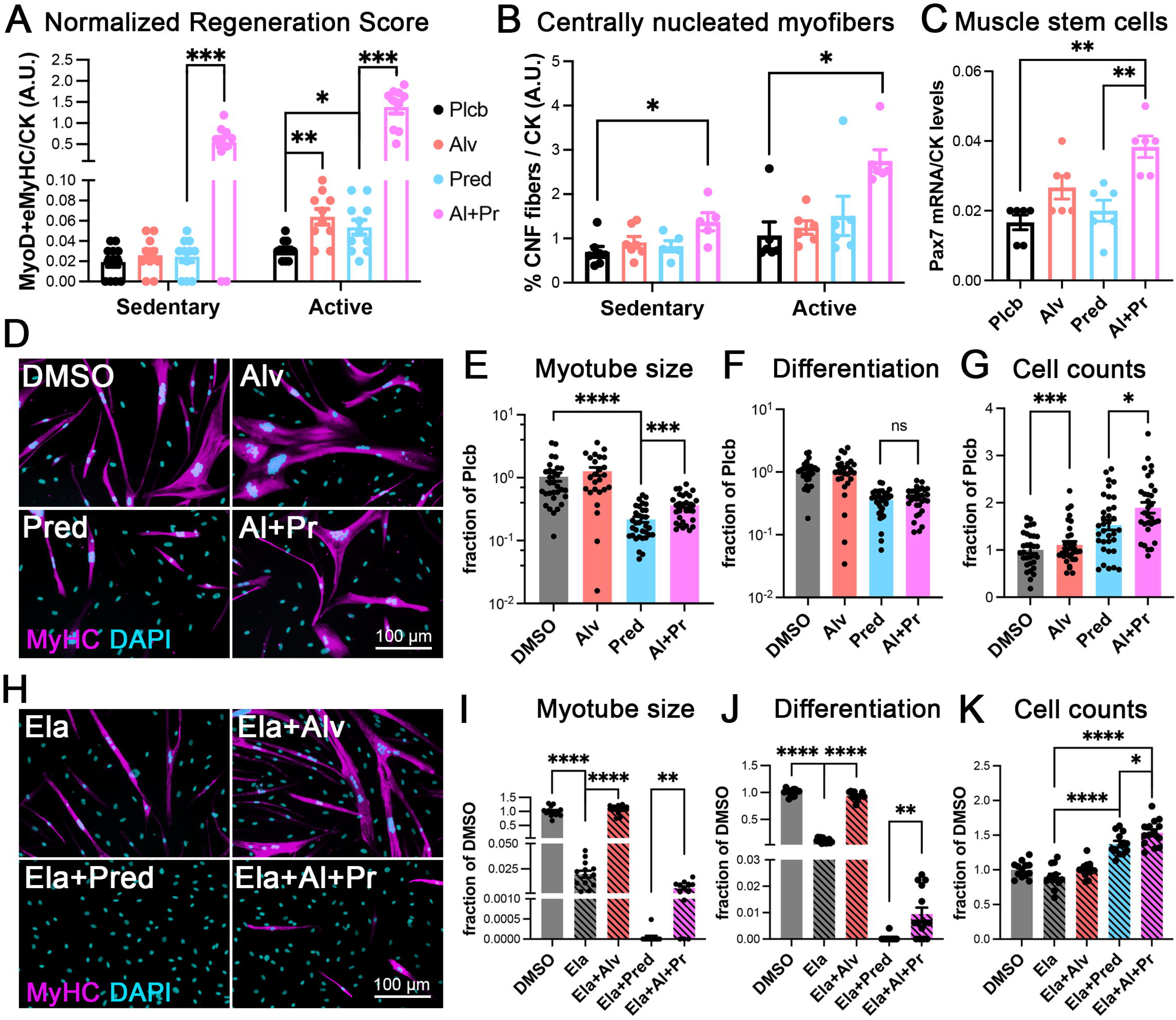
Alvelestat enhances myogenesis *in vivo*, and *in vitro* in a cell autonomous manner. **A)** Muscle extracts from mice treated with either placebo (Plcb, black), Alvelestat alone (Alv, pink), Prednisolone alone (Pred, blue) or Alvelestat + Prednisolone (Al+Pr, magenta) were homogenized and processed to detect MyoD1 and eMyHC levels via western blotting. The Normalized Regeneration Score was calculated as described in the *Results* section and plotted for each animal (individual points) sorted based on whether belonging to the sedentary or the active arm. The averages per cohort are depicted as bar height and the error bars are S.E.M. **B)** The percentage of centrally-nucleated fibers (CNF) for each animal treated as in (A) was measured using SMASH(39), then for each animal the CNF value was normalized to the average CK value for that cohort and plotted (individual points) sorted based on whether belonging to the sedentary or the active arm. The averages per cohort are depicted as bar height and the error bars are S.E.M. N = 5 at least per cohort. **C)** Pax7 expression was quantified via RNA- sequencing as a proxy for muscle stem cell abundance. N= 6 per cohort. **D-G)** Myoblasts derived from a patient with DMD were expanded and differentiated as described in the *Methods* section in the presence of either vehicle (DMSO, gray), Alvelestat (Alv, pink), Prednisolone (Pred, blue) or Alvelestat+Prednisolone (Al+Pr, magenta). Representative images of differentiated cells are shown in (D), quantification of myotube size is shown in (E), differentiation index in (F) and cell numbers in (G). **H-K)** Same cells as in D-G), cultured as in D-G) but with the addition of exogenous human ELANE. Representative images of differentiated cells are shown in (H), quantification of myotube size is shown in (I), differentiation in (J) and cell numbers in (K). * = p<0.05, ** = p<0.01, *** = p<0.001, in the indicated comparisons.

We hypothesized that Prednisolone and Alvelestat could directly affect myoblast proliferation and differentiation, in a cell-autonomous manner. To test this hypothesis, we cultured human myoblasts derived from a patient affected by DMD(11) with Prednisolone and Alvelestat, either alone or in combination, and either in the presence or absence of exogenous neutrophil elastase (Fig. 9D). While Prednisolone dramatically decreased myogenesis in these conditions, the addition of Alvelestat slightly rescued the defect caused by Prednisolone (Fig. 9D-E), by primarily promoting myoblast proliferation, both alone and in combination with Prednisolone (Fig. 9F-G). This was consistent with the increase in Pax7 mRNA observed *in vivo*, as described above (Fig. 9C). We then further pushed the system to mimic the *in vivo* scenario of high doses of ELANE in the proximity of fiber damage, which is also where myoblast migrate to, and we repeated the same experiments in the presence of additional human ELANE added exogenously. Under these conditions, ELANE decreased myogenesis (compare Ela in Fig. 9H with DMSO in Fig. 9D) as previously reported(10), and the effect was completely rescued by co-administration of Alvelestat (Fig. 9H-I). Treatment with Prednisolone in the presence of elastase completely abrogated myogenesis *in vitro* and was only slightly, but significantly, rescued by the co-administration of Alvelestat (Fig. 9H-I). An identical trend was observed when myoblast differentiation was measured, with elastase and Prednisolone decreasing it and Alvelestat increasing/rescuing it (Fig. 9J). In contrast, the effects of Prednisolone and Alvelestat on proliferation/survival in the presence of exogenous ELANE were similar to those observed when no exogenous ELANE was added (compare Fig. 9G with 9K) and as predicted by our previous experiment (Fig. 2C-D).

## DISCUSSION

The pathogenesis of Duchenne muscular dystrophy is highly complex. While gene therapy for DMD has been developed, and methods to improve it are currently being explored, it is likely that combinatorial therapies addressing also other aspects of DMD pathogenesis will eventually need to be devised. Here we show that a combination of the current mainstream standard of care for DMD, the corticosteroid Prednisolone, with the orally bioavailable neutrophil elastase inhibitor Alvelestat, yields a powerful therapeutic approach, where the potency of Prednisolone is enhanced by almost three times in the key functional outcome measures, and its most relevant side effect on muscle mass, atrophy, is reduced. It is noteworthy to mention that in patients affected by DMD neutrophil elastase expression, which is already chronically higher compared with healthy controls (24), is further induced upon treatment with standard of care corticosteroids (25, 26), further pointing to the co-administration of a corticosteroid and a neutrophil elastase inhibitor as a promising therapeutic approach.

Upon investigation of the molecular and cellular mechanisms underlying the effects of Alvelestat addition to Prednisolone, we identified the following: (i) enhancement of regenerative potential and preservation of muscle mass; (ii) increased resistance to myofiber fatigue without losing specific force; (iii) a shift of the balance between pro- and anti-inflammatory immune response towards a more pro-regenerative, anti-inflammatory state.

The positive contribution of Alvelestat to muscle regeneration is consistent with our previous finding that ELANE accumulation in dystrophic mice is associated with regeneration decline, and that *ex vivo* elastase causes myoblast death, likely via anoikis caused by elastase proteolytic activity on cell adhesion complexes (10). Indeed, myoblast survival is one of the assays we have employed here to assess *ex vivo* potency of the three neutrophil elastase inhibitors considered for this study. *In vivo* we observe a likely expansion of the muscle stem cell population, preservation of muscle mass and enhanced regeneration, in response to coadministration of Alvelestat and Prednisolone compared with both placebo and Prednisolone alone. This is consistent with the *ex vivo* data showing increased myoblast expansion and differentiation, leading to increased myotube size, when the drugs were administered together to DMD myoblasts, under culture conditions that mimic the DMD muscle microenvironment *in vivo*, where levels of ELANE are high (10). Additionally, we observed preservation of myotube size when the two drugs were added together on already formed myotubes, suggesting that overall, the preservation of muscle mass was ensured by both enhancement of regeneration and prevention of Prednisolone-induced atrophy (27). None of these effects of Alvelestat seemed to be taking place directly at the transcriptional level. Therefore, it is likely that Alvelestat only blocked elastase activity on ECM remodeling, as well as activation levels and/or integrity of cytokines, growth factors and their receptors (28–34).

The increase in resistance to fatigue could also be the result of two factors. In the quadriceps muscle group, we found a small myofiber type switch towards slower types at the expense of faster types. While the quadriceps are not the best muscles to assess myofiber switch, since the type IIb fibers (expressing Myh4) are vastly dominant in quadriceps, this trend, which goes in the same direction as the resistance to fatigue, strongly supports the notion that myofiber type switch is at least in part responsible. It must be noted that embryonic myosin heavy chain is also increased in muscles of mice treated with Alvelestat and Prednisolone, and the embryonic isoform is a slow type of muscle heavy chain, further contributing to the myofiber type switch. Another contributor to increased resistance to fatigue could be the simple fact that muscle of mice treated with Alvelestat and Prednisolone are larger than muscle treated with Prednisolone alone.

Lastly, we observed an interesting, if paradoxical, effect of the co-administration of Alvelestat and Prednisolone on the immune response and fibrosis. Overall, the combinatorial approach produced an enhanced anti-inflammatory effect by both reducing the overall immune cell burden while increasing the fraction of anti-inflammatory cells, namely T-regs, and this when compared with both placebo and Prednisolone alone. It must be noted that the macrophage burden and TGF-β levels correlated, as expected since macrophages are the main source of TGF-β in damaged muscle. The interesting and paradoxical effect is the one on fibrosis measured by picrosirius red staining: while Prednisolone alone presents significant anti-inflammatory activity compared with placebo, as expected, it is also associated with increased picrosirius red staining, which is fully rescued by co-administration with Alvelestat, even though Alvelestat alone appears to increase both inflammation and fibrosis compared with placebo. One way to explain these data is the direct effect of Alvelestat on extracellular matrix remodeling. ELANE cleaves several ECM proteins, including fibrillar and non-fibrillar collagen, laminin and elastin (10, 28) and pro-TGF-β (34). Inhibition of neutrophil elastase would then lead to a complex effect on ECM remodeling by at least directly decreasing ECM degradation and TGF-β activation. Additionally, improved regeneration and decreased necrosis would also indirectly contribute to ECM remodeling by altering the amount and nature of the immune response, and therefore the amount and nature of pro- and anti-fibrotic cytokines and other soluble factors. Another potential contributor to the increase in fibrosis observed with Prednisolone treatment is myofiber atrophy, which even alone inevitably leads to a decrease in the myofiber/ECM ratio.

In conclusion, our data show a way to enhance by approximately three times the efficacy of the most commonly prescribed, cheapest, and mutation-independent drug for DMD, Prednisolone, while decreasing some of its side effects. Additionally, it should be noted that Alvelestat has already been tested safe in clinical trials in a pediatric population (12), and is currently in phase III for anti-trypsin deficiency, thus making it a very appealing drug with a potential for quick repurpose.

## MATERIALS and METHODS

### Enzymatic activity assays

Elastase activity was measured as recommended by the manufacturer instructions for each neutrophil elastase used: Sigma for the human ELANE (Catalog number E-8140) and R&D Systems for the mouse ELANE (Catalog number 4517-SE-010). Enzymatic assays were carried out both using the buffer suggested by the manufacturer, in DMEM and in OptiMem to verify that the inhibitors behaved similarly under optimal *in vitro* conditions for each enzyme (with the manufacturer suggested buffer) and under the conditions that were going to be used in the *ex vivo* assays (in DMEM and OptiMem). In all cases the inhibitory power of each inhibitor on each enzyme was comparable across all buffers used, regardless of the absolute activities measured.

### Cell culture

All cell cultures were maintained in a humidified incubator, at 37 °C, in the presence of 5% CO_2_ and atmospheric O_2_.

Primary DMD myoblasts from the MyoLine platform(11) kindly donated by Dr. Vincent Mouly at the Institute of Myology, Paris) were maintained in growth medium (64% DMEM + 16% M199 + 20% fetal bovine serum + 25 μg/mL fetuin + 5 ng/mL hEGF + 0.5 ng/mL FGF2 + 5 μg/mL insulin + 0.2 μg/mL dexamethasone + 50 μg/mL gentamycin). To study the effect of Alvelestat (AZD9668, from either Med Chem Express or Enzo Lab, where each batch from either manufacturer was re-assessed for purity in house by NMR), BAY-678 (Sigma), and SSR69071 (Tocris) on human ELANE (Sigma) at preventing human DMD myoblast death, cells were seeded on 12 multi-well plates in growth medium. Upon reaching ∼70% confluence cells were switched to OptiMem supplemented with varying concentrations of the above-mentioned drugs, then fixed and analyzed 24 hours later. Cell counts were obtained by scoring and averaging 10 independent fields per well stained with DAPI.

Primary mouse myoblasts were isolated and cultured as previously described(35). Briefly, hindlimb muscles were dissected and immediately minced to obtain a fine slurry that was then centrifuged and filtered through a 40 μm cell strainer prior to being pre-plated on gelatin-coated plates in primary mouse myoblast growth medium (F12 + 0.4 mM CaCl_2_ + 15% horse serum + 1% penicillin/streptomycin). Two hours later the supernatant was collected and replated on gelatin-coated plates in primary mouse myoblast growth medium supplemented with 5 nM FGF2. The medium was refreshed every 2 days and cells passaged when reaching ∼70% confluence. To study the effect of Alvelestat, BAY-678, and SSR69071 on mouse ELANE (R&D Systems) at preventing primary *mdx^4cv^* myoblast death, cells were seeded on 12 multi-well plates in growth medium. Upon reaching ∼60% confluence, cells were switched to OptiMem supplemented with varying concentrations of the above-mentioned drugs, then fixed and analyzed 24 hours later. Cell counts were obtained by scoring and averaging 10 independent fields per well stained with DAPI.

To study the effect of Alvelestat, Prednisolone (Sigma) and human ELANE on DMD myoblast differentiation, cells were seeded on 12 multi-well plates coated with gelatin in growth medium. Upon reaching ∼90% confluence, cells were switched to OptiMem + 10 μg/mL insulin + 50 μg/mL gentamycin supplemented with varying concentrations of the above-mentioned drugs, then fixed and analyzed by immunofluorescence 3-4 days later. The differentiation index was measured as the percentage of all total cells that expressed myosin heavy chain (MF20 clone). Myotube area was measured with a bespoke written macro in ImageJ/Fiji as described before(10).

To study the effect of human ELANE, Alvelestat and Prednisolone (Sigma) on human DMD myotube size, cells were seeded on 12 multi-well plates coated with gelatin in growth medium. Upon reaching ∼90% confluence, cells were switched to OptiMem + 10 μg/mL insulin + 50 μg/mL gentamycin and let fully differentiate for 3-4 days prior to applying the above-mentioned drugs at varying concentrations. Myotubes were then fixed and analyzed by immunofluorescence 2 days later. Myotube area was measured with a bespoke written macro in ImageJ/Fiji as described before(36).

### Mice

*Mdx^4cv^* mice, generated on the C57Bl/6 background, were obtained from Jackson Laboratories, housed in a pathogen-free facility at the University of Liverpool, UK and used in accordance with the Animals (Scientific Procedures) Act 1986 and the EU Directive 2010/63/EU and after local ethical review and approval by Liverpool University’s Animal Welfare and Ethical Review Body (AWERB). Age-, sex- and background-matched wild type (C57Bl/6) controls were purchased from Charles River UK and housed for at least 2 weeks in the same facility before use.

To maximize randomization, twelve breeding pairs were set up at a time so that a large number of pups could be produced almost synchronously and the pups from several litters mixed and randomized at weaning. All animals enrolled in the study were randomly assigned to four groups which were blinded to the researchers carrying out the experiments, sample processing and analysis (FKJ, KAJ, LG and RE).

### Medicated diets and treatments

Diets incorporating three different doses (3, 10 or 30 mg/Kg/day) of either Alvelestat, BAY-678, SSR69071, or 1 mg/kg/day of Prednisolone or 10 mg/Kg/day Alvelestat + 1 mg/Kg/day Prednisolone were custom developed and prepared by Inotiv (formerly Envigo). Each diet was formulated to contain the same proportion of macro-nutrients as the normal chow used in the animal facility at the University of Liverpool, and a standard amount of vitamins. Each diet also contained sucralose to enhance palatability and counteract potential bitterness conferred by the drugs. A placebo diet with the same formulation as the medicated diets, and also containing sucralose but none of the drugs, was developed and obtained from Inotiv. A pilot experiment testing the palatability of each diet was conducted for one week and no differences in the amount of diet consumed by each cohort was observed. The pilot experiment also verified that the average daily consumption was in the expected range, and therefore, it was determined that mice were receiving the intended doses of each drug.

### Drug quantification in muscle and plasma

*Mdx^4cv^* mice were divided in 10 groups, each group received one of three doses of each ELANE inhibitor (3, 10 or 30 mg/Kg/day) or 1 mg/kg/day of Prednisolone in the diet for 1 week. At the end of the treatment, mice were euthanized, all at the same time of the day and their tibialis anterior and diaphragm muscles dissected after blood collection in heparinated tubes via cardiac puncture. Blood was immediately processed for plasma separation. Drug quantification via mass spectrometry was carried out by Alderley Analytical, Macclesfield, UK following standard procedures.

### Four-limb latency to fall test

To assess overall muscle fitness, mice were placed on a plastic-coated metal grid (overall size 40 x 25 cm, with cells of 1 x 1 cm), let acclimate for 30 seconds, then the grid was gently flipped upside down over a soft mat at a distance of 25 cm. The time between flipping and fall was measured using a stopwatch and the operation repeated three times per session, for a total of 2 sessions per week, at week −1 (one week before treatment started), week zero (the week when treatment started) and then weeks 3, 6 and 9 after treatment started.

To calculate the fatigue index, at each of the two days the duration of the third attempt was divided by the duration of the first attempt, representing the loss of force on that day. For each week the loss of force recorded on each of the two days assayed were averaged to yield the fatigue index in that week.

### Activity wheels – voluntary exercise assay

Mice were individually housed in cages fitted with activity wheels (Starr Life Sciences Corp) for three weeks and their activity monitored using a dedicated software that recorded the number of wheel spins per minute over the entire time. While data were acquired and recorded for all three weeks, only measurements from the last 18 days were included and considered for statistical analysis, as the first three days were deemed necessary for acclimation to the presence of the wheel in the cage.

### Creatine kinase activity assay

Creatine kinase activity assay was performed according to manufacturer instructions (Abcam) on 4 μL of serum samples obtained by centrifugation at 1,800 xg for 10 minutes of blood collected from each euthanized mouse via cardiac puncture, after 30 minutes to allow for coagulation. A standard curve was used for each plate to allow for comparison across plates. Upon plotting the kinetics over 1 hour of incubation with the substrate, it was determined that the reading at 10 minutes was faithfully within the linear part of the kinetic curve and used across all plates.

### Immunofluorescence

Tissue samples were dissected, embedded in OCT and immediately frozen on liquid nitrogen-cooled isopentane. For all immunofluorescence staining except with anti-eMyHC, sections were fixed with 4% paraformaldehyde (PFA) in PBS for 10 minutes at room temperature. After three washes in PBS, sections were permeabilized with PBS + 0.2% TritonX100 (PBST) followed by a blocking step with 3% bovine serum albumin (BSA) prior to overnight incubation with primary antibodies at 4 °C in PBS + 1% BSA. For anti-eMyHC antibody staining, sections were not fixed and a blocking solution containing non-fat milk, BSA and goat serum was used instead. Secondary antibody incubation was carried out for 1 h at room temperature in 5% horse serum, followed by 1 wash in PBST, 1 wash in PBS and counterstained with DAPI 2 μg/mL (Life Technologies), 2 washes in PBS and mounting with Vectashield (Vector Lab). For immunofluorescence of cells, human DMD myoblasts cultured and treated as described above were fixed with 4% PFA for 10 minutes at room temperature, then washed 3 times with PBS, permeabilized 10 minutes at room temperature with PBST and blocked for 1 h at room temperature with 10% horse serum. Primary antibody incubation was carried out overnight at 4 °C in PBS + 1% horse serum and then the same protocol as above was followed for secondary antibody incubation. The antibodies used were: rabbit anti-eMyHC (clone F1.652, DSHB), rat anti-laminin alpha-2 (*Sigma*) at 1:1,000, Rabbit anti-laminin (*Sigma*) at 1:1,000, mouse anti-myosin heavy chain (MF20 clone, *DSHB*) at 1:200. Secondary antibodies made in donkey and conjugated with Alexa 555, Alexa 488 and Alexa 647 (*Molecular Probes*) were used at 1:500 dilution.

Visualization of necrotic fibers was based on the observation that membrane leakage leads to accumulation of mouse IgGs inside the necrotic fibers and, therefore, detected by immunostaining with an Alexa555-labeled anti-mouse IgG antibody in combination with rabbit anti-Laminin and DAPI staining as mentioned above.

### Pathology assessment

Liver and triceps muscle samples were fixed in formalin for 48 hours, dehydrated in increasing concentrations of ethanol, before being embedded in paraffin, sectioned at 4µm thickness and stained with hematoxylin and eosin to assess tissue pathology. A board-certified veterinary pathologist scored the sections from 12 mice per cohort and assigned the scores as follows.

- Muscle pathology. Scored 1-4 for each of the following categories: Hypercontracted and necrotic fibers, regeneration, inflammatory foci, mineralization. A higher score corresponded to increased signs of pathology. The cumulative score was the sum of the scores attained by each muscle in each category. The average of the two triceps was then plotted and used for statistical analysis.
- Liver pathology. Scored 1-4 according to the presence of: extramedullary hematopoiesis (1), hydropic degeneration and hepatocellular swelling (2), inclusions (3), lobular cellularity (4).

### Force measurements

Strips of diaphragm muscle, including the central tendon and ribs were dissected and immersed in a bath of oxygenated Ringer’s solution maintained at 37 °C with the tendon on one side and the rib on the other side connected to a force transducer. After 10 minutes of acclimation, the muscle strips were gradually stretched and the twitch force measured at each length using the software Picoscope 6, until the optimal length was reached. At this point a 30-seconds long tetanic stimulation was produced and the force generated measured. After the first tetanic stimulation, muscles were rested for 5 minutes before another 20-seconds long tetanic stimulation was applied. In total 6 tetanic stimulations of 20 seconds each were applied, each separated from the next by 5 minutes rest. At the end of the measurement, the upon tendon and rib holding the diaphragm strips were removed and the length and weight of the strip were measured. Two strips per animal were assayed and the average force for each animal was plotted and used for statistical analysis. To calculate the specific force, the absolute force measured as difference between the peak measurement and the baseline measurement, was divided by the cross-sectional area, in turn calculated with the formula:

Cross-sectional area of the diaphragm strip = (Length/4.7) / (Weight/1.06)

### Western blotting

Quadriceps from *mdx^4cv^* mice treated with placebo, or the various drugs, were homogenized in RIPA buffer (150 mM NaCl, 50 mM Tris-HCl, pH 7.5, 1.0% IGEPAL, 0.1% SDS, 0.5% sodium deoxycholate) supplemented with protease inhibitor cocktail (Complete, *Roche*) and phosphatase inhibitors (1 mM Na_3_VO_4_ + 1 mM NaF) using VelociRuptor homogenizer (Scientific Laboratory Supplies) followed by incubation on ice for 30 minutes and then cleared by centrifugation at 17,000 xg for 10 min at 4 °C. Western blotting was performed as previously described(37). The antibodies used were: mouse anti- MyoD1 (BD Biosciences, clone 5.8A) at 1:500, mouse anti-eMyHC (F1.652 clone, DSHB) at 1:1000, rabbit anti-CD45 (Cell Signaling Technology) at 1:1000, rabbit anti-F4/80 (Cell Signaling Technology) at 1:1000, rat anti FOXP-3 (Thermo Scientific) at 1:500. Secondary anti-mouse, anti-rat and anti-rabbit antibodies, conjugated to horse radish peroxidase were all from Cell Signaling Technology and all used ay 1:5000. Enhanced chemiluminescence (ECL) was produced on the probed membranes using an ECL kit from Amersham.

#### Densitometry analysis

ECL images were acquired on an Amersham Imager 680 at non-saturating exposure times and band intensity quantified using the *Gel Analysis* function in ImageJ/Fiji.

### Active TGF-β ELISA assay

Muscle samples were homogenized in RIPA buffer and processed to detect the levels of active TGF-β using an ELISA kit from Novus Biologicals, according to manufacturer instructions. A standard curve was introduced in each plate to allow for normalization and comparisons across plates. Each sample was run in triplicate and the average of the three technical replicates for each animal was then used to calculate the average and standard deviation of the mean for each cohort, which were plotted and used for statistical analysis.

### Picrosirius red staining and quantification

Cryosectioned, unfixed muscles were processed as previously described(38). Briefly, sections were fixed for 1 h at 56 °C in Bouin’s fixative, washed in water, stained for 1 h in Master*Tech Picro Sirius Red, washed in 0.5 % acetic acid, dehydrated, equilibrated with xylene, and mounted using Permount™. The fluorescence emitted by Picrosirius red staining was visualized in the RFP channel of an EVOS M5000 – 10 random field from each section were imaged and for each image the percentage of collagen positive area relative to total section area in the image was measured by selecting signal positive pixels in Adobe Photoshop, deleting all background pixels, converting the resulting image to a binary image, and counting positive pixels using the ImageJ Analyze Particles function. In addition, for each image the average staining intensity was measured using the dedicated function on the EVOS microscope. The fibrotic index was then calculated as percentage area multiplied by average staining intensity to provide a proxy to a 3D quantification of the collagen amount in the image. At least ten sections per mouse for at least five different mice were scored.

### Muscle morphometry

Percentage of centrally-nucleated myofibers and percentage of necrotic fibers were measured using the functions “CNF” and “Fiber type”, respectively, of the software SMASH(39). At least ten images per animal were analyzed for at least 7 animals per cohort.

### Statistical analysis

For all experiments, unless otherwise specified (e.g. for RNA-sequencing, see below) the software GraphPad was used to carry out statistical analysis. In all cases, single pairwise comparisons were analyzed with an appropriate statistical test upon testing for normality: datasets that passed the normality test were analyzed using a parametric test (usually t-test with Welch’s correction for non-equal variance). Datasets that did not pass the normality test were analyzed with a non-parametric test, usually Mann-Whitney.

### RNA extraction and sequencing

The quadriceps muscles of several animals per cohort were pulverized in liquid nitrogen using a ceramic mortar and pestle, and then the RNA extracted using Trizol, according to manufacturer instructions. RNA quality was checked on a Bioanalyzer and the top six samples for each cohort (with RIN > 7.5) where shipped to Novogene for library preparation and sequencing, described in detail below.

#### Library Construction, Quality Control and Sequencing

Messenger RNA was purified from total RNA using poly-T oligo-attached magnetic beads. After fragmentation, the first strand cDNA was synthesized using random hexamer primers followed by the second strand cDNA synthesis. The library was ready after end repair, A-tailing, adapter ligation, size selection, amplification, and purification. The library was checked with Qubit and real-time PCR for quantification and bioanalyzer for size distribution detection. Quantified libraries will be pooled and sequenced on Illumina platforms, according to effective library concentration and data amount.

#### Clustering and sequencing

The clustering of the index-coded samples was performed according to the manufacturer’s instructions. After cluster generation, the library preparations were sequenced on an Illumina platform and paired-end reads were generated.

#### Quality control

Raw data (raw reads) of fastq format were firstly processed through in-house perl scripts. In this step, clean data (clean reads) were obtained by removing reads containing adapter, reads containing ploy-N and low-quality reads from raw data. At the same time, Q20, Q30 and GC content the clean data were calculated.

### Differential gene expression analysis

Downstream analysis was performed using a combination of programs including STAR, HTseq, Cufflink and our wrapped scripts. Alignments were parsed using STAR program and differential expressions were determined through DESeq2/edgeR. GO and KEGG enrichment were implemented by the ClusterProfiler. Gene fusion and difference of alternative splicing event were detected by Star-fusion and rMATS software.

#### Reads mapping to the reference genome

Reference genome and gene model annotation files were downloaded from genome website browser (NCBI/UCSC/Ensembl) directly. Indexes of the reference genome was built using STAR and paired-end clean reads were aligned to the reference genome using STAR (v2.5). STAR used the method of Maximal Mappable Prefix (MMP) which can generate a precise mapping result for junction reads.

#### Quantification of gene expression level

STAR will count the number of reads per gene while mapping. The counts coincide with those produced by htseq-count with default parameters. Then, the FPKM of each gene was calculated based on the length of the gene and reads count mapped to this gene. FPKM, Reads Per Kilobase of exon model per Million mapped reads, considers the effect of sequencing depth and gene length for the reads count at the same time, and is currently the most commonly used method for estimating gene expression levels (Mortazavi et al., 2008).

#### Differential expression analysis

Differential expression analysis between two conditions/groups (six biological replicates per condition) was performed using the DESeq2 R package (1.14.1). DESeq2 provide statistical routines for determining differential expression in digital gene expression data using a model based on the negative binomial distribution. The resulting P-values were adjusted using the Benjamini and Hochberg’s approach for controlling the False Discovery Rate(FDR). Genes with an adjusted P-value <0.05 found by DESeq2 were assigned as differentially expressed.

## ACKNOWLEDGMENTS

This works was supported by funding to AP from Duchenne UK, Joining Jack, Charlie’s Fund, Muscular Dystrophy Association USA and Medical Research Council UK. We wish to thank the Animal Facility at the University of Liverpool for their superb care of our mouse colonies.

## Notes

### Competing Interest Statement

The authors have declared no competing interest.

### Summary of Updates

slightly updated text (improved figure captions), and figures (changed the color palette of immunofluorescence images to be color blind-friendly. Lastly, a few typos have also been corrected.

