## Supplementary Figures for "Enhancement of Prednisolone efficacy and safety in Duchenne muscular dystrophy via neutrophil elastase inhibition"

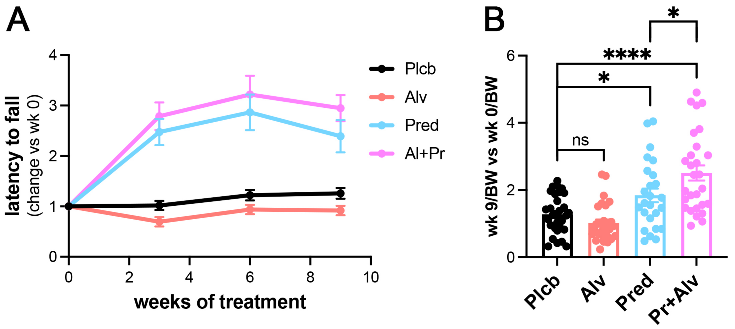


**<u>Figure S1</u>. A)** Fold change in latency to fall over 9 weeks of treatment versus week 0 were averaged across all the animals in each cohort and plotted as average +/- S.E.M. Placebo (Plcb, black), Alvelestat alone (Alv, pink), Prednisolone alone (Pred, blue) or Alvelestat + Prednisolone (Al+Pr, magenta). **B)** Fold change in latency to fall at 9 weeks versus week 0 normalized by body weight.

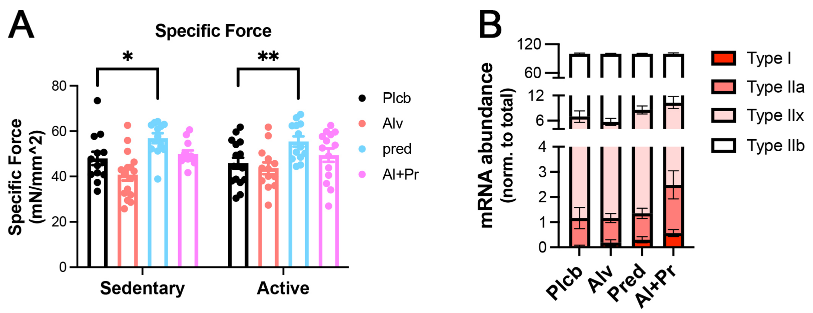


**<u>Figure S2</u>. A)** All animals treated with either placebo (Plcb, black), Alvelestat alone (Alv, pink), Prednisolone alone (Pred, blue) or Alvelestat + Prednisolone (Al+Pr, magenta) were eithanized, their diaphragm muscles dissected with the rib cage and the central tendon intact then for each animal the force elicited by a diaphragm strip (from tendon to rib) was measured and divided by the cross-sectional area of the strip as described in the Methods section to determine the tetanic specific force. N = 12 at least per cohort. **B)** Expression levels of each of the main 4 types of myosin heavy chain isoforms were measured by RNA-sequencing and plotted for each cohort, for each isoform, as a fraction of the total of all isoforms. N = 6 per cohort. * = $p<0.05$, ** = $p<0.01$, for each comparison as indicated.

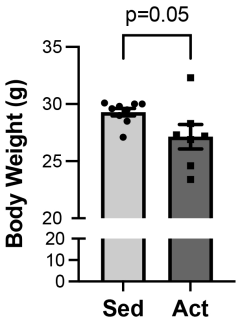


**<u>Figure S3</u>**. Changes in body weight of wild type mice that were either left sedentary or exposed to a voluntary activity wheel for three weeks.

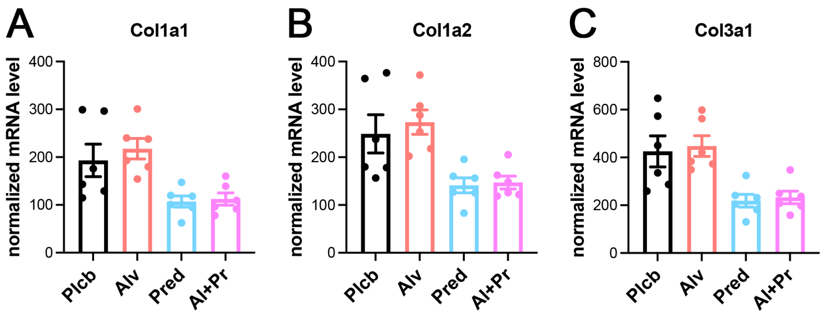


**Figure S4**. Expression levels of each of the three collagen chains that were detected in these muscle samples and that are known to form fibrillar collagen were measured by RNA-sequencing and plotted for each animal as individual points. The averages per cohort are depicted as bar height and the error bars are S.E.M. N = 6 per cohort.

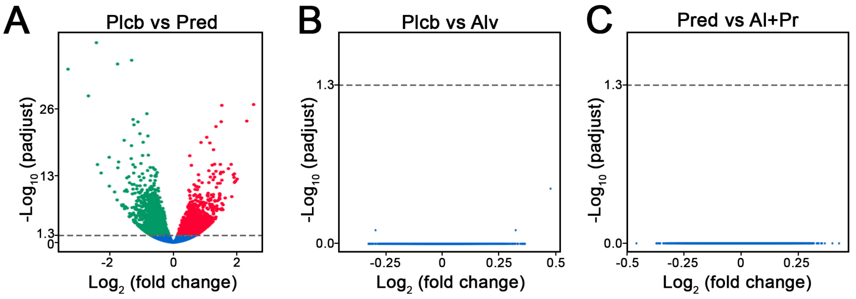


**Figure S5**. Bulk RNA-sequencing of muscles from mice treated with placebo, Alvelestat, Prednisolone or Prednisolone and Alvelestat together, was followed by differential gene expression analysis between placebo and Prednisolone (A), placebo and Alvelestat (EB), Prednisolone and Alvelestat+Prednisolone (C).
